# Structural determinants in the *Staphylococcus aureus* derived phenol-soluble modulin α2 peptide required for neutrophil formyl peptide receptor activation

**DOI:** 10.1101/2021.10.26.465826

**Authors:** Moa Viklund, Johanna Fredriksson, André Holdfeldt, Simon Lind, Henrik Franzyk, Claes Dahlgren, Martina Sundqvist, Huamei Forsman

**Author notes:** Corresponding author: Claes Dahlgren Department of Rheumatology and Inflammation Research Guldhedsgatan 10 A, S-413 46 Gothenburg, Sweden. The two authors contributed equally (Moa Viklund, Johanna Fredriksson).

## Abstract

Highly pathogenic *Staphylococcus aureus* strains produce phenol-soluble modulins (PSMs), peptides which are formylated N-terminally. Nanomolar concentrations of PSMα2 are recognized by formyl peptide receptor 2 (FPR2), but unlike the prototypic FPR2 agonist WKYMVM, PSMα2 is a biased signaling agonist. A shortened N-terminal PSMα2 variant, consisting of the five N-terminal residues, is selectively recognized by the closely related FPR1, showing that the C-terminal part of PSMα2 confers FPR2 selectivity, while the N-terminal part may interact with the FPR1 binding site. In the present study, a combined pharmacological and genetic approach, involving primary neutrophils and engineered FPR “knock-in” and “knock-out” cells, was used to gain molecular insights into FPR1 and FPR2 recognition of formyl peptides and the receptor downstream signaling induced by these peptides. In comparison to the full-length PSMα2, we show that the peptide in which the N-terminal part of PSMα2 was replaced by fMIFL (an FPR1- selective peptide agonist) potently activates both FPRs for production of superoxide anions and β- arrestin recruitment. A shortened analogue of PSMα2 (PSMα2_1-12_), lacking the nine C-terminal residues activated both FPR1 and FPR2 to produce ROS, whereas β-arrestin recruitment was only mediated through FPR1. However, a single amino acid replacement (Gly-2 to Ile-2) in PSMα2_1-12_ was sufficient to alter FPR2 signaling to include β-arrestin recruitment, highlighting a key role of Gly-2 in conferring FPR2 biased signaling. In conclusion, we provide novel structural insights into FPR1 and FPR2 recognition as well as the signaling induced by interaction with formyl peptides derived from PSMα2, originating from *Staphylococcus aureus* bacteria.

## INTRODUCTION

Neutrophils constitute the first line of our host defense against invading microbes and tissue injury, and they rely on detection of pathogen-associated molecular patterns (PAMPs) and danger-associated molecular patterns (DAMPs) produced by bacteria and/or damaged host tissues (1, 2). Peptides containing an N-formyl methionine residue is such a molecular pattern, characteristic for bacterial and mitochondrial proteins/peptides, since their biosynthesis is initiated with an N-formylated methionine (3). The specific recognition of these “non-self” peptides is achieved by their high-affinity interaction with the two formyl peptide receptors (FPR1 and FPR2), which are most abundantly expressed by human neutrophils (3, 4). The molecular structures that determine receptor selectivity are not known in detail, however, it is clear that both neutrophil FPRs recognize N-formyl peptides of different size and amino acid composition, peptides that elicit NADPH-oxidase activation to produce reactive oxygen species (ROS) (3). The essential role of FPRs in host defense has been clearly illustrated by the fact that FPR-deficient mice are more susceptible to bacterial infections than wild-type animals (5, 6).

*Staphylococcus aureus* is a major opportunistic human pathogen that causes diseases ranging from minor skin infections to severe organ damage (7, 8). Successful immune evasion by *S. aureus* is accomplished via a complex interplay involving interactions of bacterial immunomodulatory compounds and numerous virulence factors with components of the host defense systems (9, 10). Among these, fMIFL and phenol-soluble modulin α (PSMα) are such N-formylated peptides that are preferentially recognized by FPR1 and FPR2, respectively (4, 11–14). PSMα peptides secreted by community-associated methicillin-resistant *S.aureus* (CA-MRSA) strains have dual effects, since they in higher concentrations are cytotoxic and lyse both viable and apoptotic neutrophils, while at nanomolar concentrations they act as selective FPR2 agonists, stimulating pro-inflammatory responses (15–17). In addition, PSMα peptides contribute to biofilm development and trigger rapid formation of neutrophil extracellular traps (NETs) (18, 19). Hence, neutrophil FPR activation by PSMα peptides undoubtedly contributes to the outcome being either a successful elimination of bacteria by the innate immune system of the host or establishment of a persistent infection leading to tissue inflammation.

The formyl peptide-sensing FPRs belong to the large family of G protein-coupled receptors (GPCRs), and they share many of the signaling pathways and regulatory mechanisms typically associated with this receptor class. Recently, we showed that PSMα peptides are biased FPR2 agonists, which similarly to many FPR2-interacting lipopeptides induce: (i) a transient rise in the cytosolic concentration of Ca^2+^, (ii) ERK1/2 phosphorylation, and (iii) an activation of the ROS-producing enzyme system, whereas they, (iv) lack the ability to induce recruitment of *β*-arrestin (13, 20, 21). This is in line with the emerging concept of biased signaling and functional selectivity described for many GPCRs, also including FPR1, which can adopt multiple activation states and mediate biased signaling, i.e., in favor of one or another among multiple downstream signaling pathways (22, 23). It could be hypothesized, that the ability to produce biased signaling FPR2 agonists that affect neutrophil functions, has been evolved by *S. aureus* bacteria to avoid host defense activities (24). Therefore, understanding of FPR recognition and downstream signaling upon interaction with formyl peptides is highly relevant when designing novel FPR-based immunomodulatory therapeutics. The recently published structure of FPR2, crystalized in its active conformation complexed with the high-affinity agonist WKYMVm (a dual agonist for both FPRs, but with a preference for FPR2), revealed the residues critical for high affinity in ligand binding (25, 26). The crystal structure of FPR1 is currently not available, but studies with computational docking and receptor mutagenesis have suggested that the N-terminal fMet residue reaches deep into the ligand-binding pocket, and that high-affinity binding most likely involves multiple non-contiguous residues that must be properly positioned via the folding of all extracellular and transmembrane receptor domains (25). Earlier studies have suggested that the size of the proposed binding pocket, available for FPR1, is limited to peptides with four or at the most five amino acid residues (27). Nevertheless, our earlier studies strongly suggest that FPR preference is not determined solely by the amino acids which are directly inserted into the presumed agonist binding pocket of the receptor, but also by peptide parts not directly in contact with this pocket (14).

In the present study, we aimed to gain knowledge on FPR recognition of formyl peptides and ligand-directed FPR signaling, leading to ROS production and β-arrestin recruitment. We performed structure-activity relationship (SAR) analysis with a number of variants of the PSMα2 peptide in primary neutrophils and in genetically modified FPR “knock-in” and “knock-out” cells in order to compare their ability to trigger one or the other of the signaling pathways. Our data reveal critical roles of both the N-terminal part (in particular the second residue) and the C-terminal part of PSMα2 in determining receptor selectivity and FPR2 biased agonism. These structural insights into how PSMα peptides interact with one or both FPRs in innate immune cells to evoke either balanced or biased signaling may assist design of new therapeutics for treating *S. aureus* infections by inducing desirable inflammatory responses.

## MATERIALS AND METHODS

### Ethics statement

This study, conducted at the Sahlgrenska Academy in Sweden, includes blood from buffy coats of healthy individuals obtained from the blood bank at Sahlgrenska University Hospital, Gothenburg, Sweden. According to the Swedish legislation section code 4§ 3p SFS 2003:460 (Lag om etikprövning av forskning som avser människor), no ethical approval was needed since the buffy coats were provided anonymously and could not be traced back to a specific donor.

### Reagents and peptide synthesis

Dextran and Ficoll-Paque were obtained from GE-Healthcare (Waukesha, WI, USA). Isoluminol was purchased from Sigma-Aldrich and horse radish peroxidase (HRP) was from Roche Diagnostics (Bromma, Sweden). The hexapeptides WKYMVM/m were purchased from Alta Bioscience (University of Birmingham, Birmingham, United Kingdom), while formyl- mitocryptide COX1 was a kind gift from Hidehito Mukai (Nagahama Institute of Bio-Science, Japan). A sample of PSMα2 (fMGIIAGIIKVIKSLIEQFTGK) was obtained from EMC microcollection GmbH (Tuebingen, Germany), while another sample and all PSMα2 analogues were synthesized in house using procedures previously reported (28). All peptide stocks were made in DMSO and further dilutions were made in Krebs-Ringer phosphate buffer containing glucose (10 mM), Ca^2+^ (1 mM) and Mg^2+^ (1.2 mM) (KRG, pH 7.3). Cyclosporin H (CysH) was kindly provided by Novartis Pharma (Basel, Switzerland). The FPR2-specific peptidomimetic antagonist (CN6: RhB-[Lys-βNPhe]_6_-NH_2_; where RhB = Rhodamine B and βNPhe = N- phenylmethyl-β-alanine) was synthesized as described (29). Ficoll-Paque was obtained from Amersham Biosciences. RPMI 1640, fetal calf serum (FCS), penicillin/streptomycin (PEST) and G418 were from PAA Laboratories GmbH, Austria.

### FPR-mediated β-arrestin recruitment assay

The ability of FPR2 agonists to promote β-arrestin recruitment was evaluated in the PathHunter® eXpress system with CHO-K1 FPR cells and accompanying reagents from DiscoverX (Fremont, USA), as described (21, 30). In brief, CHO cells overexpressing FPR1 or FPR2 were diluted in Cell Plating Reagent before seeded in tissue culture treated 96-well plates (10.000 cells/well) and incubated for approximately 20 h (37°C, 5% CO_2_). Cells were then stimulated with agonists for 90 min at 37°C followed by addition of detection solution for 60 min after which chemiluminescence was measured using a CLARIOstar platereader (BMG Labtech, Germany).

### Isolation of blood-derived neutrophils and culture of FPR1- and FPR2-deficient neutrophil-like HL60 cells

Blood neutrophils were isolated as described by Böyum (31, 32) using buffy coats from healthy volunteers. After dextran sedimentation at 1 × g, hypotonic lysis of the remaining erythrocytes, the neutrophils obtained by centrifugation in a Ficoll-Paque gradient were washed twice in KRG. The cells were resuspended in KRG (1 × 10^7^/ml) and stored on ice until use.

The HL60 cells were cultured in RPMI-1640 medium supplemented with 10% FBS, 1% Penicillin-Streptomycin and 2 mM L-glutamine at 37°C, 5% CO_2_. For differentiation into neutrophil-like cells, DMSO (1%) was added to cell culture and the cells were allowed to differentiate 6 days. On the day of experiments, cells were washed once and then resuspended in KRG, and stored on ice until use.

FPR1- and FPR2-deficient HL60 lines were created as described (33). In brief, PCR amplicons were designed to amplify and sequence the CRISPR targeted regions out of genomic DNA isolated from HL60 cells. CRISPR gRNAs were designed and oligos synthesized, annealed, and cloned into the all-in-one Cas9-2A-GFP/sgRNA vector pXPR004 (BROAD Institute). NEON electroporation (THERMO) was used to deliver the Cas9/sgRNA constructs into HL60 cells and clones were expanded. Cells were differentiated in 1% DMSO for 6 days into neutrophil-like cells and used for ROS production experiments.

### Measurement of superoxide anion production

The production of superoxide anion by the neutrophil NADPH-oxidase was measured by isoluminol-amplified chemiluminescence (CL) in a six-channel Biolumat LB 9505 (Berthold Co, Wildbad, Germany) as described earlier (34). In short neutrophils or neutrophil-like HL60 cells were mixed (in a total volume of 900 μl containing 1 – 2.5 × 10^5^ cells) with HRP (4 U/ml), and isoluminol (6 × 10^-5^ M) in KRG, pre-incubated at 37°C after which the stimulus (100 μl) was added. The light emission was measured continuously. When required, the specific receptor antagonists were included in the CL mixture for 5 min at 37°C before stimulation.

### Data collection and statistical analysis

Data analysis was performed using GraphPad Prism 9 (Graphpad Software, San Diego, CA, USA). EC_50_ values for PSMα2 analogues were determined by fitting a sigmoidal dose-response curve with variable slope to data normalized to the maximal response (100%) using the “log (agonist) versus normalized response” in GraphPad Prism. The statistical analysis was performed on raw data-values by using a two-tailed paired Student’s *t*-test or a repeated measures one-way ANOVA followed by Dunnett’s multiple comparison test. Statistical significance is indicated by \**p* < 0.05.

## RESULTS

### Dissection of the involvement of FPR1 and FPR2 in neutrophil recognition of peptide agonists – a pharmacological approach

Neutrophils have evolved FPR1 and FPR2 for the recognition of peptides containing an fMet group at the N-terminus (4). These peptides are a hallmark of bacterial and mitochondrial protein synthesis, and they are recognized by the innate immune system as pathogen-associated molecular patterns (PAMPs) and danger-associated molecular patterns (DAMPs) (3). A common pharmacological approach for determining receptor preference for a formyl peptide is to apply receptor antagonists with known selectivity for either FPR1 or FPR2, respectively (35). This is clearly illustrated by the effects of the potent FPR1-selective antagonist CysH that inhibits NADPH-oxidase generation of ROS in human neutrophils upon stimulation with the FPR1- selective agonist fMLF (Fig. 1A and D). Expectedly, CysH had no effect on ROS release, when triggered by the FPR2-selective peptide agonist WKYMVM (Fig 1B and D). In contrast, the most potent FPR2-selective antagonist, the peptidomimetic CN6 (i.e., RhB-[Lys-βNPhe]_6_-NH_2_), designed by combining structural elements of two potent FPR2 antagonists (i.e., PBP_10_ and Pam- [Lys-βNspe]_6_-NH_2_ (29)), inhibited the WKYMVM- but not the fMLF-induced response (Fig 1A,B and D). As opposed to the inhibition profiles observed for fMLF and WKYMVM, the two antagonists had very small effects on responses induced by the N-terminal fragment of the mitochondrial COX1 protein or the hexapeptide WKYMVm with a D-Met as C-terminus (Fig 1C and D). A complete inhibition of the activation induced by either of these agonists could be achieved only when the two antagonists were applied in combination (Fig 1C, D), clearly showing that COX1 as well as WKYMVm are recognized by both FPRs when expressed in human neutrophils.

**Figure 1.**
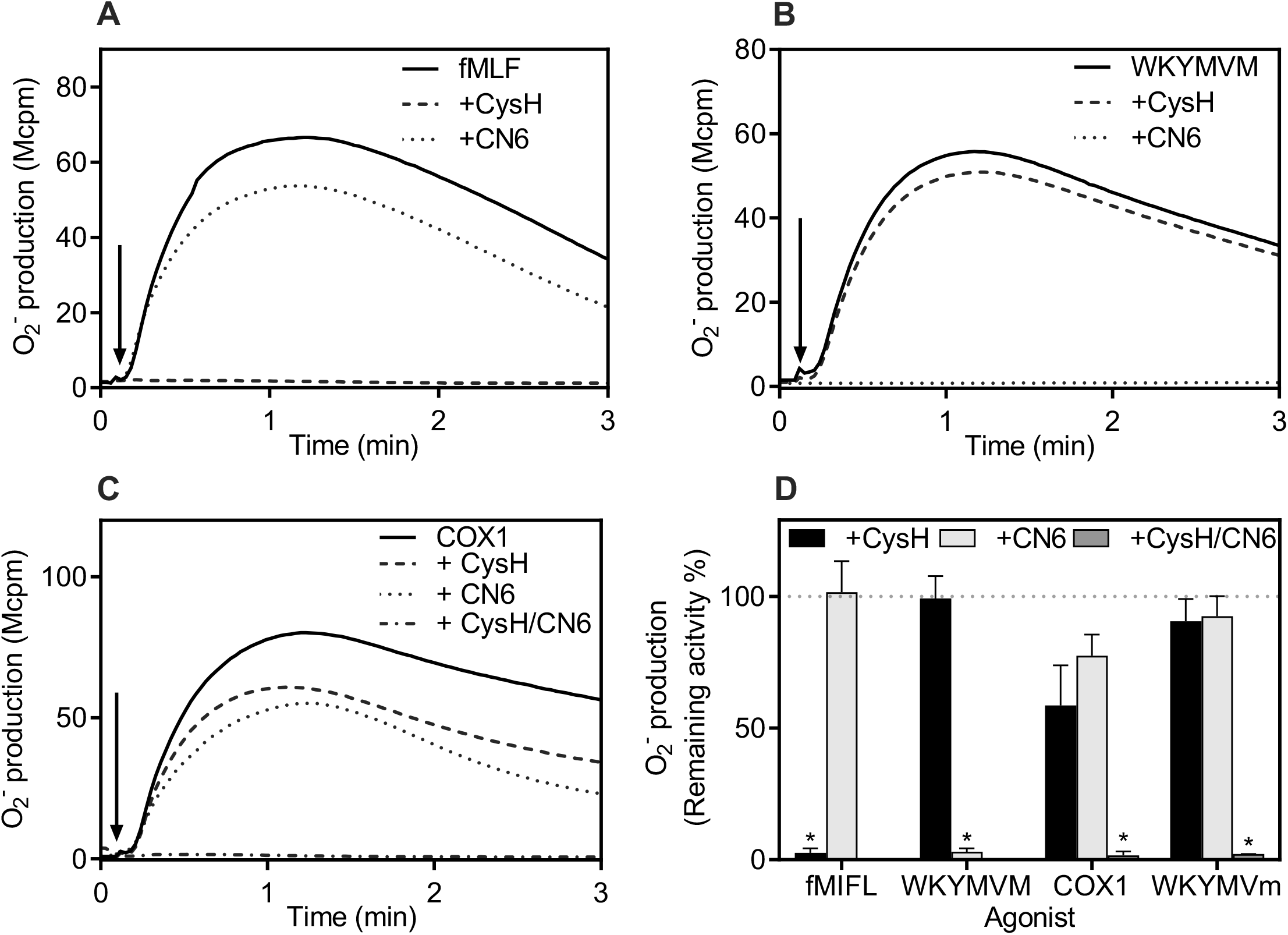
Determination of FPR-selective recognition of neutrophil-activating agonists using well-characterized receptor-selective antagonists. Activation of the superoxide anion (O_2_^-^)-generating neutrophil NADPH-oxidase was measured. **A-C)** Neutrophils (10^5^ cells) were incubated for 5 min at 37°C in the absence (no additive, solid lines) or presence of FPR-selective antagonist(s). An activating agonist was added as indicated by the arrows. The antagonists used were CysH (FPR1-selective; 1 μM, dashed lines) and CN6 (FPR2-selective; 100 nM; dotted lines). **A)** The FPR1-selective agonist was fMLF (100 nM); **B)** The FPR2-selective agonist was WKYMVM (100 nM); C) The dual FPR1/FPR2 agonist was COX1 (100 nM). One representative experiment out of three independent experiments is shown. **D)** Summary of the inhibitory effects of FPR antagonists on the response induced by 100 nM of fMLF, WKYMVM, or the two dual agonists COX1 and WKYMVm. Data are presented as percent remaining O_2_^-^ activity for each agonist in the presence of either one or both antagonists, while 100% is the agonist response in the absence of any antagonists (n=3, mean + SEM).

### Genetic “knock-in” and “knock-out” approaches in dissection of the engagement of FPR1 and FPR2

An alternative to the pharmacological characterization described above, is to use a genetic approach to determine receptor preference for a formyl peptide. In the present study, we applied both FPR “knock-in” and “knock-out” strategies to overexpress and abolish individual FPRs, respectively. For the “knock-out” strategy, human neutrophils cannot be used due to the fact that these cells are very short-lived, and therefore very difficult to manipulate genetically. Instead, the CRISPR-Cas9 technique was applied to HL60 cells to generate FPR knock-out (FPR^-/-^) neutrophil- like cells after DMSO differentiation (33). Using these neutrophil-like cells, we show that the known FPR1-selective agonist fMLF activated the FPR2^-/-^ cells, but not the FPR1^-/-^ cells to produce ROS (Fig 2A, B). In contrast, the FPR1^-/-^ cells, but not the FPR2^-/-^ cells, were activated by the FPR2-selective agonist WKYMVM (Fig 2A, B). The kinetics of ROS production from these genetically modified FPR^-/-^ neutrophil-like cells was very similar to that obtained from primary human neutrophils (Fig 1), i.e., with a rapid onset and termination within five minutes. Also, and in agreement with the receptor profile obtained using the pharmacological approach with human neutrophils (Fig 1), the dualism of COX1 and WKYMVm was confirmed; both these peptides activated FPR1^-/-^ as well as FPR2^-/-^ cells (Fig 2C). For the “knock-in” strategy we used CHO cells (that lack endogenous expression of the two FPRs) overexpressing FPRs that were co-expressed with β-arrestin, which allowed us to study FPR downstream β-arrestin recruitment as described previously (21). By using this system, we confirmed the receptor preference for the FPR1-selective agonist fMLF, which initiated signals that promoted recruitment of β-arrestin in FPR1 cells, but not in FPR2-overexpressing cells (Fig 2D). In contrast, the FPR2-selective agonist WKYMVM induced β-arrestin recruitment solely in FPR2 cells, but not in FPR1-overexpressing cells, while the dual FPR agonist COX1 induced β-arrestin recruitment in both cell types (Fig 2D).

**Figure 2.**
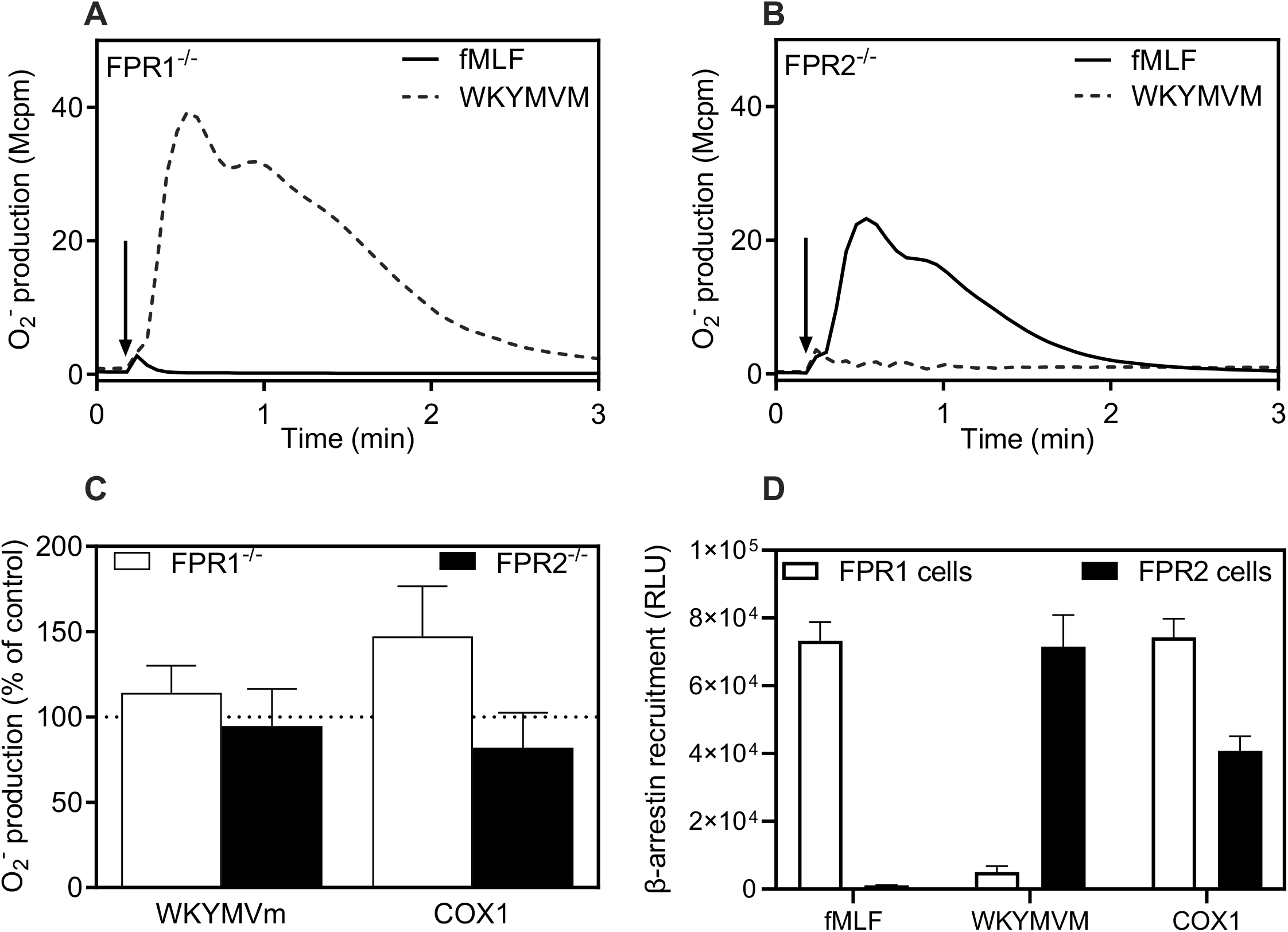
FPR selectivity as determined by genetic knock-in and knock-out approaches. **A-B)** FPR agonist-induced O_2_^-^ release from DMSO-differentiated neutrophil-like HL60 cells deficient in either FPR1 (i.e., FPR1^-/-^, **A**) or FPR2 (i.e., FPR2^-/-^, **B**) was measured. Cells were stimulated with fMLF (100 nM, solid lines) or WKYMVM (100 nM, dashed lines). One representative experiment out of three with different passages of HL60 cells is shown. **C)** The NADPH-oxidase activation induced by the dual FPR agonists WKYMVm (100 nM) and COX1 (100 nM) in FPR1^-/-^ cells (white bars) and FPR2^-/-^ cells (black bars). Data are presented as percent of the control responses, induced by WKYMVM (100 nM) for FPR1^-/-^ cells and by fMLF (100 nM) for FPR2^-/-^ cells (mean + SEM, n=3. **D)** The ability of the FPR1-selective agonist fMLF (10 nM), the FPR2-selective agonist WKYMVM (25 nM), and the dual agonist COX1 (100 nM) to recruit β-arrestin was measured in CHO cells overexpressing FPR1 (white bars) or FPR2 (black bars). The bar graphs show the *β*-galactosidase activity (relative light units, RLU), reflecting the degree of β-arrestin recruitment (mean + SEM, n=3).

Collectively, these data clearly show that we with the genetic approach confirm receptor selectivity for the most commonly used FPR-targeting pharmacological tool compounds, and the cells used, thus, constitute excellent models for in-depth mechanistic studies.

### Both FPR1 and FPR2 recognize N-formyl peptides in which the C-terminal part plays a critical role in determining receptor preference

The precise structural features that determine whether a formyl peptide is selectively recognized by FPR1 and/or FPR2 remain largely unknown, except for the findings that the fMet residue plays an essential role for recognition and that the size of the peptide is important (3, 4). *S. aureus* releases several formyl peptides including the short fMIFL and the 21-residue PSMα2 (Table 1A), which are selectively recognized by FPR1 and FPR2, respectively (4). In agreement with this, we show that neutrophils were activated by PSMα2 through FPR2, as illustrated by a complete inhibition by the FPR2-selective antagonist CN6, but not by the FPR1-selective antagonist CysH in the neutrophil ROS release system (Fig 3A). Despite the FPR2 preference for the longer PSMα2, the receptor preference for the 5-residue N-terminal fragment, PSMα2_1-5_ (i.e., fMGIIA) was changed. Similar to fMIFL (the other *S. aureus*-produced FPR1 agonist) PSMα2_1-5_ (Table 1A) selectively activated FPR1 with respect to neutrophil superoxide release, but only at much higher (100-fold) concentrations (Fig 3B, C). The receptor preferences, determined by applying the pharmacological inhibition approach, were fully supported by data obtained with the genetically modified HL60 cells deficient in either FPR1 or FPR2. Thus, FPR1 was activated by nanomolar (nM) concentrations of fMIFL and micromolar (µM) concentrations of PSMα2_1-5_, and both peptides induced ROS production in the FPR2^-/-^ cells only (Fig 3D, E). In comparison to PSMα2_1- 5_ that required µM concentrations to activate FPR2^-/-^ cells, fMIFL was more potent with a maximal response in FPR2^-/-^ cells at a concentration as low as 0.1 nM, whereas FPR1^-/-^ cells were activated by fMIFL at concentrations >50 nM (Fig 3D, E). In contrast to the activation by these two short formyl peptides, the longer PSMα2 peptide selectively activated cells that express a functional FPR2, and it lacked effect on FPR2^-/-^ cells even at concentrations ≥500 nM (Fig 3F). These data clearly show that the 5-residue N-terminal fragment of PSMα2 was selectively recognized by FPR1, and that the C-terminal part of PSMα2 plays a critical role for its preference for FPR2.

**Figure 3.**
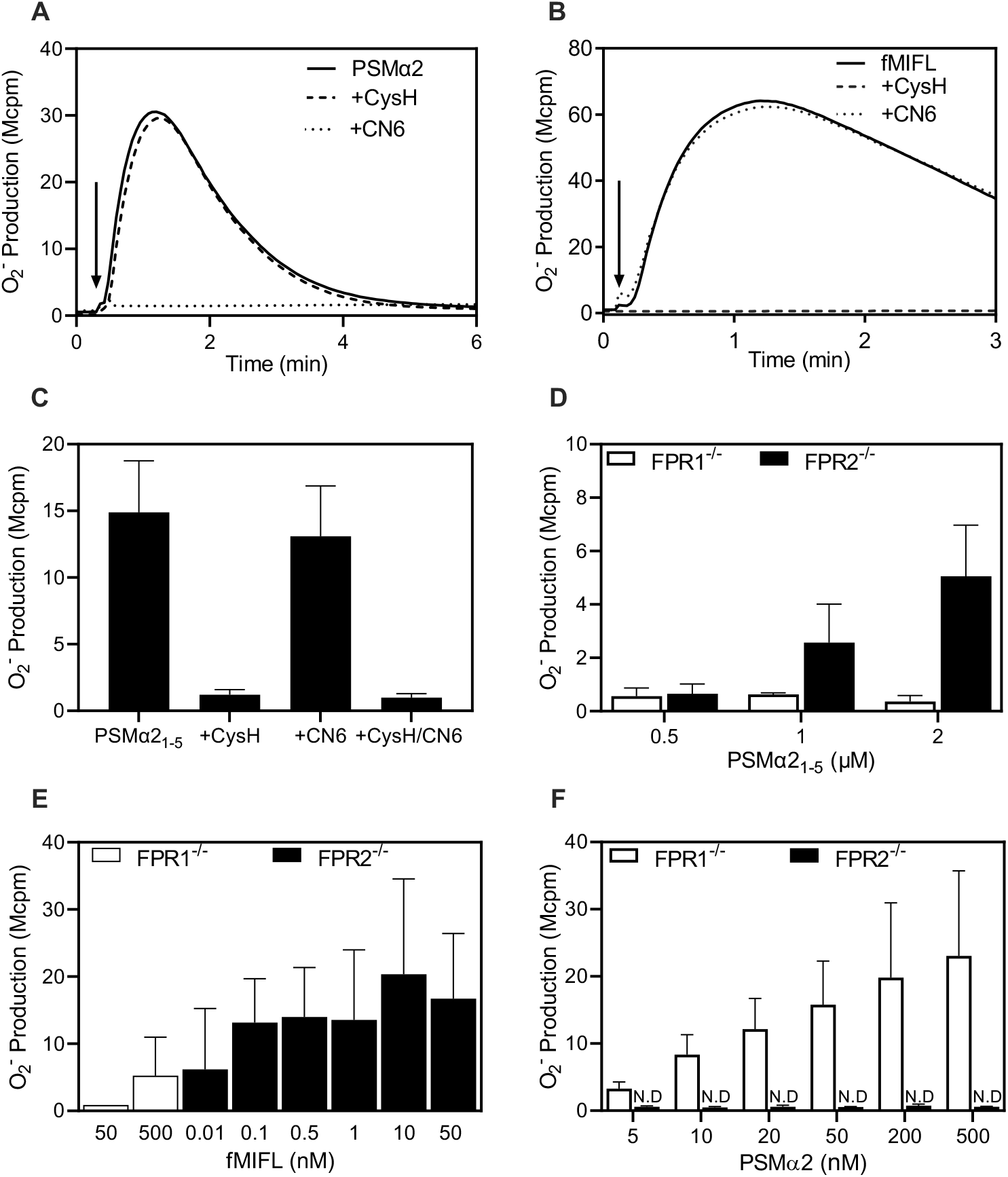
Recognition characteristics of PSMα2 and fMILF, two peptides produced by *S. aureus*. Activation of FPRs by fMIFL, PSMα2 and the N-terminal fragment PSMα2_1-5_ of the latter was measured as neutrophil O_2_^-^ production induced by the agonists. The receptor mediating the response was determined by a pharmacological approach (**A-C**) and a genetic knock-out approach (**D-F**). **A-B)** Experiments performed with PSMα2 (50 nM; **A**) and fMIFL (1 nM; **B**) as agonists. The response induced in the absence of any antagonist (solid lines), in the presence of the FPR1 antagonist (CysH; 1 µM, dashed lines) or the FPR2 antagonist (CN6; 100 nM, dotted lines) are shown. One representative experiment out of three independent experiments are shown. **C)** Peak neutrophil O_2_^-^ production was determined when induced by PSMα2_1-5_ (1 µM) alone and in presence of a FPR subtype-selective antagonist alone (CysH for FPR1; CN6 for FPR2) and when combined. **D-F**) Superoxide production induced in FPR2 expressing (FPR1^-/-^; white bars) and FPR1-expressing (FPR2^-/-^; black bars) HL60 cells by different concentrations of PSMα2_1-5_ (**D),** fMIFL **(E)** and, PSMα2 **(F)**. Data are presented as peak responses (mean + SEM, n = 3). N.D. non-detectable.

**Table 1.** **A** Peptide length and amino acid sequence of the different PSMα2 variants

| Peptide name | 1 | 2 | 3 | 4 | 5 | 6 | 7 | 8 | 9 | 10 | 11 | 12 | 13-21 |
| --- | --- | --- | --- | --- | --- | --- | --- | --- | --- | --- | --- | --- | --- |
| PSMα2<sub>1-5</sub> | fM | G | I | I | A | | | | | | | | |
| PSMα2<sub>1-12</sub> | fM | G | I | I | A | G | I | I | K | F | I | K | |
| PSMα2 | fM | G | I | I | A | G | I | I | K | F | I | K | G – K* |
| fMIFL-PSMα2<sub>5-21</sub> | fM | I | F | L | A | G | I | I | K | F | I | K | G – K* |

\*PSMα2 and fMIFL-PSMα2<sub>5-21</sub> contain the fragment GLIEKFTGK at positions 13 - 21**B**Peptide length and amino acid sequence of PSMα2<sub>1-12</sub> variants

### The two neutrophil FPRs recognize a 21-residue chimeric formyl peptide and both the N- and C-terminal parts are important

Despite the fact that fMIFL and PSMα2 are potent FPR1- and FPR2-selective agonists, respectively, the 21-residue chimeric peptide fMIFL-PSMα2_5-21_ (in which three amino acids in the N-terminal part (i.e., G^2^I^3^I^4^ of PSMα2 were replaced with IFL), retained the neutrophil activating capacity (Fig 4A, B). However, only a very low level of inhibition was achieved when either of the two FPR subtype-selective antagonists CysH or CN6 were applied alone, and an increased inhibition was obtained when the antagonists were applied together (Fig 4A, B). These data suggest that fMIFL-PSMα2_5-21_ interacts with both FPRs and, compared to PSMα2, gained affinity for FPR1 while retaining affinity for FPR2. Its neutrophil activation potency was also increased (EC_50_ *∼* 1 nM; Fig 4C), compared to that of PSMα2 (EC_50_ *∼* 50 nM, (14)). The receptor recognition profile of fMIFL-PSMα2_5-21_, as determined in the neutrophil ROS production assay, was confirmed in experiments with the “knock-out” HL60 cells, showing that the chimeric fMIFL- PSMα2_5-21_ activated both FPR^-/-^ cells, but with a higher potency for FPR2 than FPR1 (Fig 4D).

**Figure 4.**
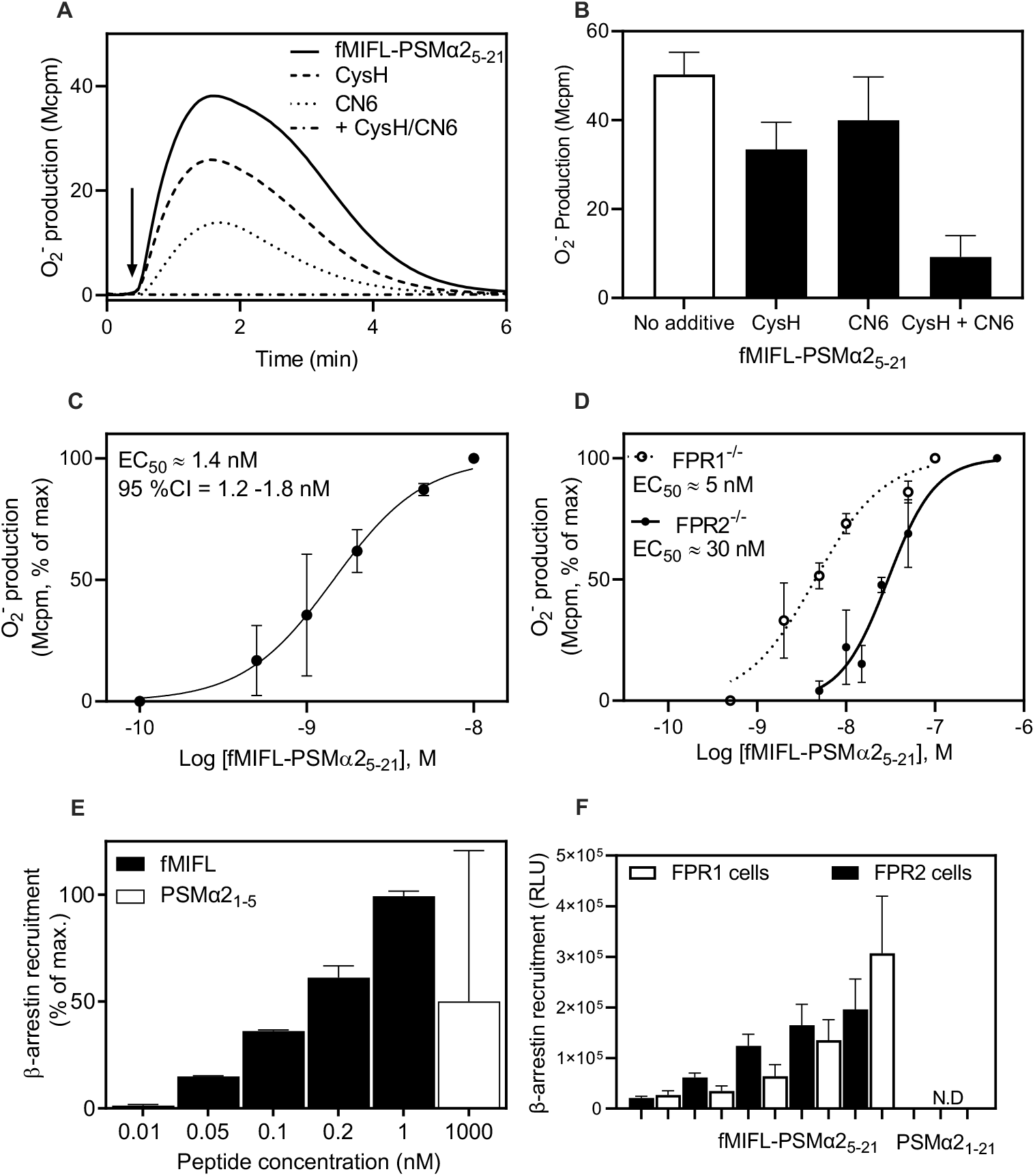
FPR recognition pattern of the chimeric fMIFL-PSMα2_5-21_ peptide. **A-B**) FPR preference for fMIFL-PSMα2_5-21_ in human neutrophils as determined through the inhibitory effects of FPR-selective antagonists. **A)** The neutrophil production of O_2_^-^ was measured when induced by fMIFL-PSMα2_5-21_ (5 nM) in the absence (solid line) or presence of an FPR1 antagonist (CycH; dashed), an FPR2 antagonist (CN6; dotted), or both antagonists in combination (dash-dotted). One representative experiment out of three independent experiments is shown. **B)** Summary of the effects of the FPR subtype-selective antagonists alone or in combination (black bars). The response induced by fMIFL-PSMα2_5-21_ in the absence of any antagonist is also shown as control (white bar); the results are presented as the peak O_2_^-^ response (mean + SEM, n = 3). **C-D)** Concentration dependent activation of the O_2_^-^ producing NADPH-oxidase (peak values) when induced by fMIFL-PSMα2_5-21_ in neutrophils **(C)** and HL60 cells overexpressing FPR2 (FPR1^-/-^; open circles in **D**) or FPR1 (FPR2^-/-^; filled circles in **D**). Data are shown with EC_50_ values determined as normalized peak responses from 3 independent experiments (mean + SEM, n = 3). **E-F)** Recruitment of β-arrestin in CHO cells overexpressing FPR1 or FPR2 when induced by different peptide agonists. **E)** β-arrestin recruitment induced by different concentrations of fMIFL in FPR1-overexpressing cells (black bars; mean + SEM, n=3). The response induced by PSMα2_1- 5_ (1 µM; white bar) is shown for comparison. Data are presented in percent of the response induced by the FPR1 agonist fMLF (100 nM). **F)** β-arrestin recruitment induced by the chimeric 21-residue fMIFL-PSMα2_5-21_ in cells overexpressing FPR1 (white bars) or FPR2 (black bars). Data represent mean + SEM, n=3. The inability (ND) of PSMα2_1-21_ (500 nM) to recruit β-arrestin is shown for comparison (ND = non-detectable).

The ability of these formyl peptides to recruit β-arrestin upon FPR recognition was further investigated by using the “knock-in” strategy (Fig 2D). In agreement with the receptor recognition data, the two short peptides fMIFL and fMGIIA promoted recruitment of β-arrestin in FPR1- overexpressing cells (Fig 4E) and the former being the more potent. When fMIFL was conjugated with the C-terminal part of PSMα2, the chimeric fMIFL-PSMα2_5-21_ peptide triggered β-arrestin recruitment by both FPRs with some preference for FPR2 over FPR1 (Fig 4F). This was in contrast to the full-length PSMα2 that lacked the ability to trigger any β-arrestin recruitment (Fig 4F). These findings infer that the biased FPR2 signaling capacity is determined by the N-terminal part of the peptide.

In summary, both the N-terminal and the C-terminal parts of the two 21-residue long peptides (PSMα2 and fMIFL-PSMα2_5-21_) play critical roles for FPR preference and downstream signaling.

### The C-terminal truncated PSMα2_1-12_ peptide activates the ROS-producing system down-stream of both FPR1 and FPR2, whereas β-arrestin is recruited only by FPR1

To investigate the structural basis for recognition of peptide agonists by FPRs in more detail, we generated a C-terminal truncated PSMα2 peptide (PSMα2_1-12_; Table 1A) and some chimeric/variant analogues of this (Table 1B). In addition to fMIFL-PSMα2_5-12_ (i.e., displaying G^2^→I^2^, I^3^→F^3^, and I^4^→L^4^ substitutions), six 12 amino acids long N-terminal peptides-variants (i.e., G^2^I^3^I^4^ adjacent to fMet) were generated and included in the study (Table 1B). Similar to PSMα2, also the shorter PSMα2_1-12_ activated neutrophils to produce ROS, but based on the antagonist inhibitory effects this peptide classifies as a dual FPR agonist since both antagonists were needed to achieve complete inhibition (Fig 5A). Interestingly, the chimeric peptide fMIFL-PSMα2_5-12_, in which the three amino acids G^2^I^3^I^4^ adjacent to fMet were replaced by I^2^F^3^L^4^, proved to be a more potent agonist but with the receptor dualism retained (Fig 5A).

**Table 1.** **B** Peptide length and amino acid sequence of PSMα2_1-12_ variants

| Peptide name | 1 | 2 | 3 | 4 | 5-12 |
| --- | --- | --- | --- | --- | --- |
| [I<sup>2</sup>]-PSMα2<sub>1-12</sub> | fM | I | I | I | A-K<sup>#</sup> |
| [F<sup>3</sup>]-PSMα2<sub>1-12</sub> | fM | G | F | I | A-K<sup>#</sup> |
| [L<sup>4</sup>]-PSMα2<sub>1-12</sub> | fM | G | I | L | A-K<sup>#</sup> |
| [I<sup>2</sup>F<sup>3</sup>]-PSMα2<sub>1-12</sub> | fM | I | F | I | A-K<sup>#</sup> |
| [I<sup>2</sup>L<sup>4</sup>]-PSMα2<sub>1-12</sub> | fM | I | I | L | A-K<sup>#</sup> |
| [F<sup>3</sup>L<sup>4</sup>]-PSMα2<sub>1-12</sub> | fM | G | F | L | A-K<sup>#</sup> |
| [I<sup>2</sup>F<sup>3</sup>L<sup>4</sup>]-PSMα2<sub>1-12</sub> | fM | I | F | L | A-K<sup>#</sup> |

#The PSMα2 variants and fMIFL-PSMα2<sub>5-21</sub> contain the fragment AGIIKFIK at positions 5-12

**Figure 5.**
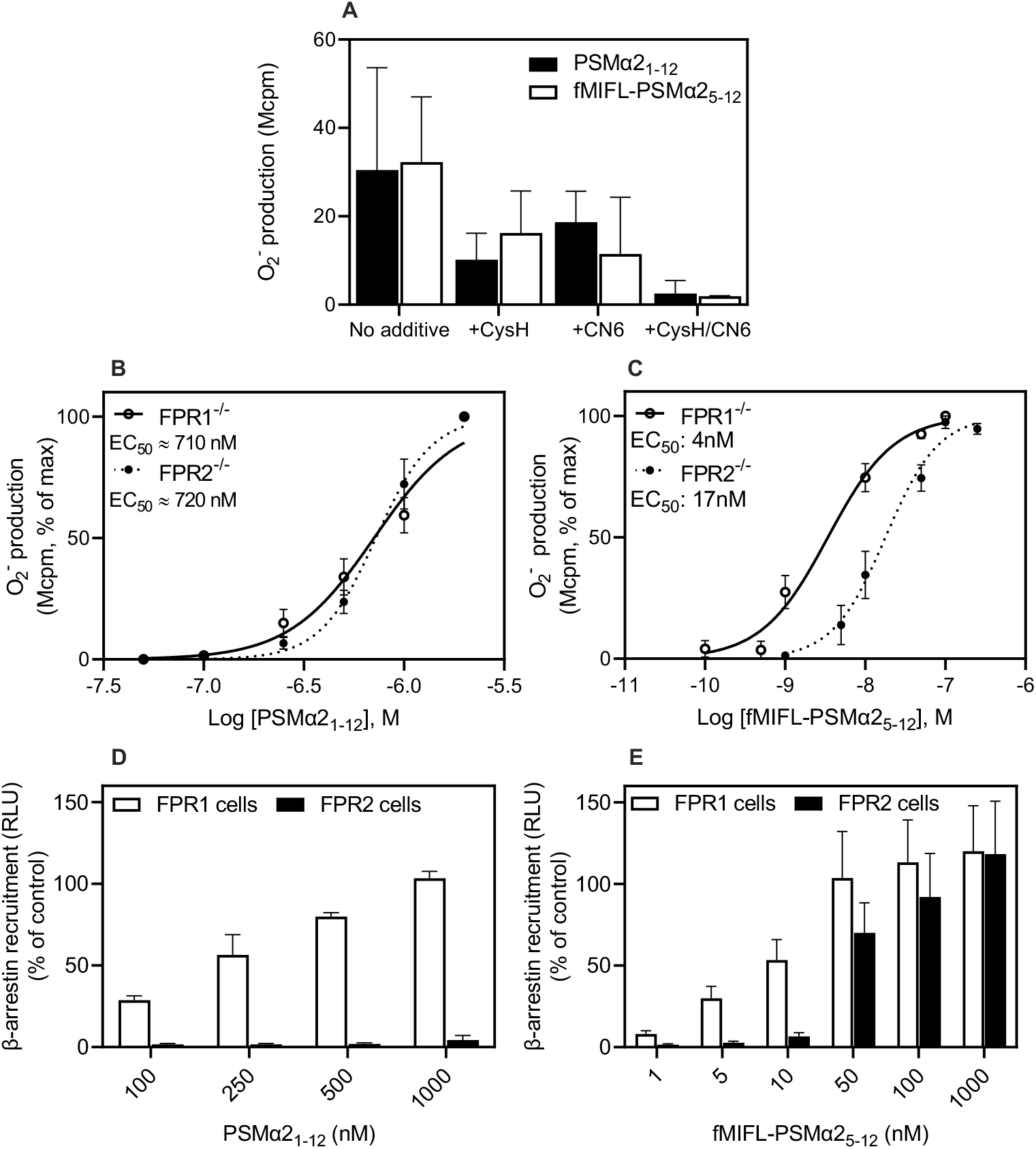
FPR recognition and signaling induced by the truncated PSMα2_1-12_ peptide and the chimeric fMIFL-PSMα2_1-12_ peptide. **A)** FPR preference for PSMα2_1-12_ and fMIFL-PSMα2_5-12_ in human neutrophils as determined by the inhibitory activity of FPR subtype-selective antagonists CysH (FPR1 selective) and CN6 (FPR2 selective). Neutrophils were activated with PSMα2_1-12_ (1 µM, black bars) or fMIFL- PSMα2_5-12_ (1 nM, white bars) in the absence or presence of an FPR selective antagonist alone or when the two were combined as indicated. Data are presented as the peak responses in the absence of any antagonist (No additive) and in the presence of one antagonist or the two combined (mean + SEM, n=3). **B-C)** Production of O_2_^-^ (peak values) induced in HL60 cells by different concentrations of PSMα2_1-12_ (**B**) or fMIFL-PSMα2_5-12_ (**C**). The activities in cells expressing FPR2 (FPR1^-/-^; open circles) or FPR1 (FPR2^-/-^; filled circles) were determined; the data are presented as normalized responses with EC_50_ values determined from 3 independent experiments (mean + SEM, n = 3). **D-E)** Recruitment of β-arrestin induced by different concentrations of PSMα2_1-12_ (**D**) or fMIFL-PSMα2_5-12_ (**E**). Data are presented as percent of the activity induced by fMLF (100nM; for FPR1-overexpressing cells) and WKYMVM (100nM; for FPR2-overexpressing cells) and represent mean + SEM (n = 3).

In agreement with the receptor profiles obtained from the experiments with human neutrophils, the dual agonistic properties of the shorter PSMα2_1-12_ and the chimeric fMIFL-PSMα2_5-12_ peptides were evident also with cells in which one of the FPRs was genetically deleted (Fig 5B and C). The potencies in activation (as measured by EC_50_ values) of PSMα2_1-12_ for both FPRs were *∼* 700 nM (596 – 838 nM for FPR1^-/-^ cells and 645 – 809 nM for FPR2^-/-^ cells with 95% CI). In comparison to PSMα2_1-12,_ the chimeric fMIFL-PSMα2_5-12_ variant was a considerably more potent agonist in both FPR1^-/-^ cells (EC_50_ *∼*5 nM; 95% CI: 4 – 7 nM) and FPR2^-/-^ cells (EC_50_ *∼*20 nM; 95% CI: 22 – 42 nM).

The ability of these two peptides to recruit β-arrestin was examined in the receptor “knock-in” system. Similar to the full-length PSMα2, also the shorter PSMα2_1-12_ lacked the ability to trigger *β*-arrestin recruitment in FPR2-overexpressing cells (Fig 5D). In contrast, but in agreement with the data showing that PSMα2_1-12_ was a dual agonist that activates also FPR1, the peptide dose- dependently induced *β*-arrestin recruitment in FPR1-overexpressing cells (Fig 5D). The chimeric fMIFL-PSMα2_5-12_ proved not only to be a potent dual FPR agonist in activation of the NADPH-oxidase with an EC_50_ in the low nanomolar range, but the peptide also recruited β-arrestin in both FPR1- and FPR2-overexpressing cells, with a higher potency for FPR1 (Fig 5E).

Taken together, these data show that the FPR2-selective agonist PSMα2 and its non-selective shorter analogue PSMα2_1-12_ transduce biased signals downstream from FPR2, leading to ROS production, but not to β-arrestin recruitment. In contrast, the chimeric fMIFL-PSMα2_5-12_ variant is a potent dual FPR agonist with respect to both induction of ROS production and *β*-arrestin recruitment.

### Amino acid substitutions in the N-terminus of the PSMα2_1-12_ peptide change the receptor activation profile as determined by FPR1- and FPR2-mediated ROS release and β-arrestin recruitment

To further elucidate how the amino acids in the N-terminal of PSMα2_1-12_ affect FPR1 and FPR2 signaling leading to ROS production and β-arrestin recruitment, we designed different PSMα2_1-12_ analogues/variants that all retained the fMet residue, but displayed substitutions in the adjacent three positions (i.e., the G^2^I^3^I^4^ triplet); the eight C-terminal residues were kept/identical in all peptide variants (Table 1B). Like the parental PSMα2_1-12_, which induce activation of the NADPH-oxidase through both FPRs (Fig 5B), all these substituted analogues activated neutrophil-like cells expressing FPRs to produce ROS (Fig 6A), albeit to different levels and receptor preference. Regarding the interaction with FPR1, all variants except the I^4^→L^4^ substitution were full ROS activation agonists and the peptides containing two of the IFL amino acids were the most potent, with EC_50_ values close to that of chimeric fMILF-PSMα2_5-12_ peptide (Fig 6A). The potency of the peptides in which one of the amino acids in PSMα2_1-12_ was changed were less potent than fMILF- PSMα2_5-12_ with the analogue displaying an I^4^→L^4^ substitution being the least potent (Fig 6A). This peptide was also the least potent agonist to recruit *β*-arrestin down-stream of FPR1. All the other peptides variants were as potent with respect to *β*-arrestin recruitment as fMILF-PSMα2_5-12_ (Fig 6C).

**Figure 6.**
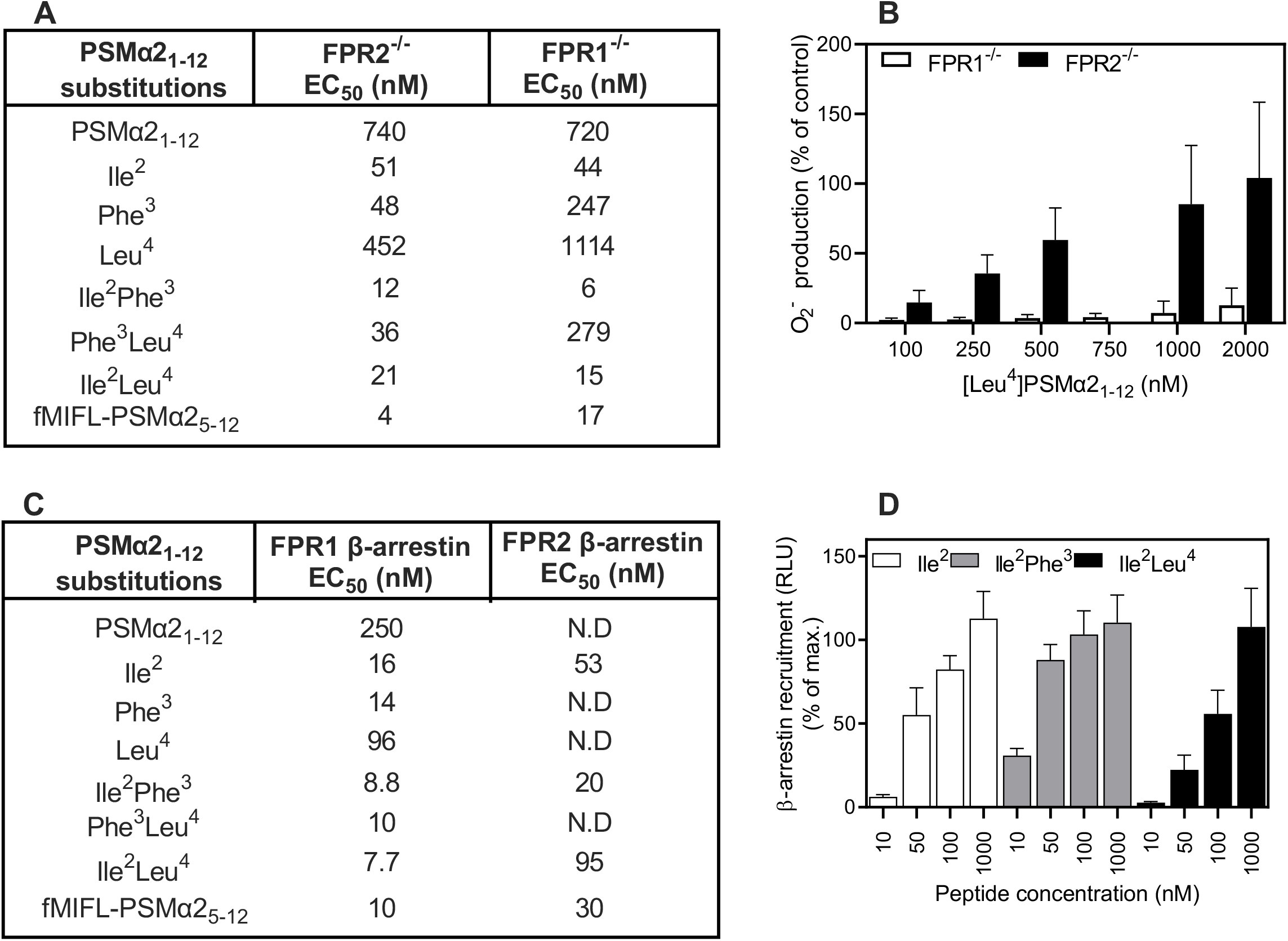
FPR recognition and signaling induced by PSMα2_1-12_ and variants with substitutions in the N-terminal part. **A)** Production of O_2_^-^ induced by PSMα2_1-12_ and variants of this peptide in FPR1- and FPR2- deficient HL60 cells shown as EC_50_ values determined for each peptide from three independent experiments. **B)** Production of O_2_^-^ in HL60 cells expressing FPR1 (FPR2^-/-^; black bars) and FPR2 (FPR1^-/-^; white bars) when induced by different concentrations of the PSMα2_1-12_ peptide variant in which I in position 4 was replaced by an L. Data are presented as percentage of the superoxide production (peak values) induced by WKYMVM (100 nM) in FPR1^-/-^ cells and fMLF (100nM) in FPR2^-/-^ cells (mean + SEM, n = 3). **C)** Recruitment of β-arrestin in cells overexpressing FPR1 or FPR2 when induced by PSMα2_1-12_ and variants of this peptide shown as EC_50_ values determined for each peptide from three independent experiments. EC_50_ was determined for each peptide (ND= non-detectable. **D)** Recruitment of β-arrestin in cells overexpressing FPR2 when induced by different concentrations of three PSMα2_1-12_ variants having a G^2^→I^2^ substitution in common. The recruitment was compared to that induced by WKYMVM (100 nM; 100%) and expressed as mean + SEM (n = 3).

Regarding the interaction with FPR2, the three peptides in which G^2^ in PSMα2_1-12_ was replaced by an Ile (i.e., G^2^→I; G^2^I^3^→IF; G^2^I^4^→IL) were all full agonists with increased potencies as compared to PSMα2_1-12_ with EC_50_ values fairly similar to that of fMILF-PSMα2_5-12_ (Fig 6A). In contrast, the three peptides lacking the Ile^2^ residue in position 2 were low (seen for I^3^→P; I^3^I^4^→PL) or very low-ROS activating (I^4^→L) peptides (Fig 6A and B). This clearly shows that many alterations close to the N-terminus confer retained ability to engage both FPRs for ROS production. The analogue with an I^4^→L^4^ substitution is FPR1-selective as almost no ROS production was induced in the FPR1^-/-^ cells even when a concentration of 2 µM was used to activate the cells (Fig 6B). This suggests that I^4^ in PSMα2_1-12_ plays a key role in determining FPR subtype selectivity. Noticeably, all analogues displaying Ile in position 2, including the peptide with only this substitution, were not only FPR1 agonists, but also induced ROS production through FPR2 (Fig 6A), suggesting that the second residue has a critical role in conferring FPR2 interaction.

Next, we examined the ability of these substituted analogues to promote *β*-arrestin recruitment downstream of activation of FPR2. The three peptides in which G^2^ in PSMα2_1-12_ was retained (I^3^→F; I^4^→L; I^2^I^4^→PL) shared the inability of the parent PSMα2_1-12_ to promote FPR2 to recruit *β*-arrestin (Fig 6C). The three peptides in which G^2^ in PSMα2_1-12_ was replaced by an I (G^2^→I; G^2^I^3^→IF; G^2^I^4^→IL) triggered, however, a dose dependent recruitment of *β*-arrestin in FPR2- overexpressing cells (Fig 6D), suggesting a critical role of Ile in position 2 for conferring an ability to trigger *β*-arrestin recruitment through FPR2 activation.

In summary, the data presented show that all PSMα2_1-12_ analogues with one or more residues (I, F and/or L) from fMIFL display an improved FPR1 activation potency in both ROS release and *β*- arrestin recruitment. Critical roles of amino acids in the N-terminal part of PSMα2_1-12_ were disclosed also for activation of FPR2.

## Discussion

In the present study, we investigated how the N- and C-terminal parts of the *S. aureus*-produced PSMα2 formyl peptide influence the recognition by FPRs and the receptor downstream signaling leading to an activation of the NADPH-oxidase and β-arrestin recruitment. Our results show that both termini of the peptide constructs used, play important roles in determining FPR2 selectivity and biased signaling, as well as for conferring an ability to activate not only FPR2 but also FPR1. These conclusions were drawn from our characterization data and comparison of several modified PSMα2 analogues, differing in length and amino acid composition of the N-terminal part (i.e., the three amino acids adjacent to the fMet residue). Overall, the present results show that both FPR1 and FPR2 recognize formyl peptides, but with different recognition and signaling profiles. The finding that FPR1 is a high-affinity receptor for formyl peptides is well-established, while there is no consensus as to whether formyl peptides are endogenous ligands also for FPR2; this is mainly due to the fact that FPR2 originally was shown to recognize non-formyl peptides as well as non-peptide agonists (3, 4).

The molecular basis for FPR recognition of formyl peptides remains elusive, albeit earlier studies have suggested that the binding pocket available in FPR1 is limited to four or at the most five residues, and that fMet is directly involved in this interaction with multiple non-contiguous residues positioned via proper folding of all extracellular and transmembrane domains in the receptor (3, 4, 25, 27). Recently, the crystal structure of FPR2 was resolved in an active signaling state (complexed with the high-affinity agonist WKYMVm), which revealed the presence of a widely open and deep ligand binding pocket created through conformational rearrangements, initiated by an outward shift of helix VI (25, 36). It should be noticed that although the co- crystalized WKYMVm peptide is recognized preferentially by FPR2, it is a dual agonist that potently activates also FPR1 (37). Thus, it is reasonable to assume that FPR1 and FPR2 may possess similar binding sites also for formyl peptides, however, with somewhat different restrictions on which specific molecular features that can fit the respective agonist binding pocket. Notably, the formyl peptides used in the above docking studies were very short, and thus only involved a limited number of molecular contacts with the receptors. Hence, the models developed from these experiments may not be suitable for prediction of how longer peptides interact with the receptors. Future approaches aiming to understand the structural basis for FPR1 and FPR2 recognition of not only short peptide agonists, but also for longer formyl peptides, may eventually provide more specific molecular details of FPR-ligand interactions.

With respect to the impact of the length of formyl peptides on FPR1 and FPR2 recognition, it seems that FPR1 preferentially interacts with shorter peptides. In contrast, FPR2 exhibits a preference for longer peptides. This is supported by the fact that in addition to the 21-residues long PSMα2, FPR2 selectively recognizes a number of longer formyl peptides arising from the N-terminal parts of mitochondrial DNA-encoded proteins, including the 15-residue MCT-2 peptide (from cytochrome *b*) and the 20-residue ND4 (from the NADH dehydrogenase subunit 4) (38). Shorter C-terminally truncated variants of these peptides are either inactive or activate solely FPR1 or both FPRs (14, 39–41). It is clear that the removal of a fairly large part of the C-terminal of PSMα2 and fMIFL-PSMα2_5-21_, affects receptor recognition and activation. The ability of the deleted part to fit to and directly interact with the receptor parts suggested to define the agonist binding pocket of the FPRs, must be fairly limited (25, 26). Earlier computer modeling and site-directed receptor mutagenesis experiments have suggested that different structural determinants exist for FPR1 and FPR2 with respect to interaction with short formyl peptides (i.e., 3 to 5 amino acids). Also the length of such peptides and the composition of the C-terminal part was found to be of importance for FPR2 but not critical for FPR1 binding (42).

In the current study, we show that in contrast to the PSMα2 peptide generated by *S.aureus* bacteria, C-terminally truncated analogues (i.e., PSMα2_1-5_ and PSMα2_1-12_) activate FPR1, albeit they were not very potent agonists (i.e., EC_50_ values >500 nM). In addition, the chimeric fMIFL-PSMα2_5-21_ was found to act as a potent dual agonist for both FPR1 and FPR2, suggesting that even though longer formyl peptides often gain in FPR2 selectivity, this does not exclude recognition by FPR1. Clearly, recognition of formyl peptides by both FPRs is not determined solely by the N-terminal 4-5 amino acids, suggested to fit directly into the binding pocket, since also the C-terminal part was found to contribute. The importance of the C-terminal part in conferring FPR subtype selectivity is in fact supported by earlier studies, showing that a peptide, in which the hydrophobic Ile^11^ positioned close to the C terminus of a dual FPR agonist was removed (or replaced with Ala), was no longer recognized by FPR2 whereas its ability to be recognized by FPR1 was retained (14). In the present study, we applied in addition to a traditional pharmacological approach, also a genetic approach by generating HL60 cells deficient in FPR1 or FPR2 (33). These model cells were used to gain insights into FPR signaling, leading to activation of the neutrophil ROS-producing NADPH-oxidase. The results obtained via this approach, regarding receptor selectivity for the different formyl peptides, essentially confirmed those obtained with subtype-selective FPR inhibitors in human neutrophils expressing both FPR1 and FPR2. The respective receptor selectivity for the two *S. aureus*-produced formyl peptides PSMα2 and fMIFL was confirmed by using these FPR-deficient cells. Thus, while the fMIFL peptide was a potent activating agonist in FPR2^-/-^ cells, these cells were non-responsive to PSMα; in contrast fMIFL was only weakly active in FPR1^-/-^ cells even at *∼* 50.000-fold higher concentrations (Fig 4). The genetic approach was particularly useful for dissecting the engagement of individual FPRs when a peptide agonist displayed affinity for both FPRs. We could thus, not only confirm that PSMα2_1-12_ activated both FPR1 and FPR2 for neutrophil ROS production with equal potency, but also clearly demonstrate that the shortened I^4^→L^4^ PSMα2_1-12_ variant no longer was an FPR1 agonist. This finding suggests that Ile in position 4 within PSMα2_1-12_ plays a critical role in enabling a FPR1-mediated response. Our results show that PSMα2 is biased FPR2 agonist not able to activate the receptor to recruit β- arrestin, whereas signals that activate the NADPH-oxidase were generated. To understand the structural basis for this biased FPR2 signaling, we designed several variants of PSMα2. We found that the C-terminally truncated PSMα2_1-12_ activated FPR2, and the PSMα2 signaling profile of the peptide was retained despite the truncation. Interestingly, when the N-terminal amino acid sequence G^2^I^3^I^4^ was replaced by I^2^F^3^L^4^ the signaling properties of new construct were changed. Thus, fMIFL-PSMα2_5-21_ and its shortened analogue, fMIFL-PSMα2_5-12_, both activated FPR2, generating signaling that activated the NADPH-oxidase activation system with concomitant recruitment of β-arrestin. In fact, a single amino acid substitution (G^2^→I^2^) was sufficient to trigger: (i) an enhanced ROS production, and (ii) FPR2 to recruit *β*-arrestin. Collectively, these findings show that the N-terminal residues in PSMα peptides play key roles in determining which signaling pathways become activated upon recognition by FPR2.

Even if the N-terminus of formyl peptides is the part recognized by the binding pocket in FPR2 (26), this interaction appears to be modulated by the C-terminal part protruding from the binding pocket. Thus, the amino acids in both ends of the peptide affect the subsequent conformational change of the receptor, and by that determine the receptor down-steam signaling profile, and a fairly small structural modification of the activating agonist is obviously sufficient to alter signaling and the functional outcome of the activation.

Other FPR2 agonists, structurally unrelated to PSMα2, have earlier been shown to trigger a biased signaling/functional selective response (20, 21, 30, 43). The concept of biased signaling applies also to FPR1 as illustrated by the results presented in a recent study, showing that signaling induced by the novel FPR1-selective small-molecule agonist RE-04-001 was strongly biased toward the PLC-PIP_2_-Ca^2+^ and the ERK1/2 activation pathways, but away from β-arrestin recruitment (44). In addition, it has also been shown that a pyridazin-3(2 H)-one-based agonists, recognized by FPR1, display a strongly biased signaling toward ERK1/2 phosphorylation and away from Ca^2+^ mobilization (45).

In conclusion, we present novel structural insights into FPR recognition of PSMα2 (a peptide produced by pathogenic *S. aureus* bacteria) and the induced downstream signaling. Our data reveal that both the N-terminal part (in particular the residue adjacent to fMet) and the C-terminal part of PSMα2 (and shortened analogues) play key roles in determining receptor selectivity and FPR2 biased agonism. The full-length PSMα2 selectively activated FPR2 to induce ROS production, while no β-arrestin recruitment was triggered. The shorter PSMα2 analogues activated both FPR1 and FPR2, but β-arrestin recruitment was only induced down-stream of activated FPR1. Finally, our results highlight that Gly in position 2 within PSMα2_1-12_ appears to be a determinant for inducing a ligand-directed FPR2 sub-active conformation that promotes β-arrestin recruitment. These molecular insights obtained with two *S. aureus*-produced formyl peptides, capable of activating the innate immune system via the FPR pattern-recognition receptors, have increased our understanding of host-pathogen molecular interactions, which may facilitate design of potential therapeutics against severe infections.

## Acknowledgments

We thank for the valuable suggestions provided by the members of the Phagocyte Research Group. The technical assistance provided by Linda Bergqvist is also acknowledged.

## Author contributions

HF oversaw all the aspects of the study. HF and CD designed the study with input from Henrik Franzyk who contributed with the synthesis work. MW, JF, AH, SL, MS performed the experiments with input from HF and CD. All co-authors contributed to the conception of the study, experimental analytical performance, and continuously discussion on the experiment design. HF wrote the original draft of the manuscript and all authors edited, revised and approved the final version submitted.

## Data and materials availability

All data needed to evaluate the conclusions are presented in the article. The raw data utilized in this article and additional data related to this article are available from the corresponding author (CD) upon reasonable request.

## Abbreviations

FPR: formyl peptide receptor
GPCR: G protein-coupled receptor
PSMα: phenol-soluble modulin α
ROS: reactive oxygen species
CysH: cyclosporine H
NADPH-oxidase: nicotinamide adenine dinucleotide phosphate oxidase
DMSO: dimethyl sulfoxide
HRP: horseradish peroxidase
KRG: Krebs-Ringer phosphate buffer

